# The presence of an ESBL-encoding plasmid reported during a *Klebsiella pneumoniae* nosocomial outbreak in the United Kingdom

**DOI:** 10.1101/2024.12.09.627553

**Authors:** Stephen Mark Edward Fordham, Anna Mantzouratou, Elizabeth Anne Sheridan

## Abstract

The presence of extended-spectrum β-lactamase (ESBL) encoding plasmids in bacteria contributes towards rising resistance rates, mortality and healthcare costs in clinical settings. An EBSL-encoding plasmid, pESBL-PH was identified during a nosocomial outbreak of *Klebsiella pneumoniae* ST628 at a United Kingdom general district hospital in 2018. A plasmid from the earliest 2018 *K. pneumoniae* strain discovered during the outbreak was assembled using both Oxford nanopore long reads and illumina short reads, yielding a fully closed plasmid, pESBL-PH-2018. pESBL-PH-2018 was queried against the complete NCBI RefSeq Plasmid Database, comprising 93,823 plasmids, downloaded on July 16, 2024. To identify structurally similar plasmids, strict thresholds were applied: both a shared hash ratio > 0.9 and a mash similarity ≥ 0.98. This returned 61 plasmids belonging to 13 unique sequence types (STs) hosts. The plasmids were detected in 13 unique countries, dating from 2012-2023, associated with a range of infections including bacteremia. Low numbers of single nucleotide polymorphisms (SNPs) were identified between pESBL-PH-2018 and query plasmids, further confirming their relatedness. The AMR region of the plasmids varied; interestingly IS*26* mediated-tandem amplification of resistance genes, including the ESBL *bla*_CTX-M-15_ was identified in two independent strains raising their copy number to three. Furthermore, the genomic background of strains carrying a pESBL-PH-2018-like plasmid were analyzed, revealing truncation of the chromosomal *ompK36* porin gene and carbapenemase resistance gene carriage on accessory plasmids in 17.85% and 26.78% of strains with a complete chromosome available. This analysis reveals the widespread dissemination of an ESBL-encoding plasmid in a background of resistance-encoding strains, requiring active surveillance.

## Introduction

The World Health Organization (WHO) recognizes both third generation cephalosporin (3GCR) and carbapenem resistant Gram-negative *Klebsiella pneumoniae* (*K. pneumoniae*) as a critical threat to human health [1]. Specifically, 3GCR *K. pneumoniae* are associated with increased in-hospital mortality and longer hospital stays, alongside increased healthcare costs [2,3]. Notably, the extended spectrum β-lactamase (ESBL) gene, *bla*_CTX-M-15_, encoding CTX-M-15 has been reported worldwide [4,5]. While several studies have considered the clonal outbreak of clinical *K. pneumoniae* strains encoding the ESBL CTX-M-15, such as ST101 [6], ST1427 [7], ST307 [8], and ST394 [9], among others, ESBL genes are usually present on large plasmids that can circulate between species. *K. pneumoniae* strains are exceptional reservoirs for AMR genes, capable of both acquiring and disseminating plasmids across members of its own population and other Enterobacteriaceae species [10].

The horizontal transfer of ESBL-encoding plasmids has been documented across several *K. pneumoniae* sequence types (STs), in addition to other Enterobacteriaceae species [11,12]. Notably, plasmid transmission involving multiple strains has been recorded. A 120-kbp IncFIIK conjugative plasmid encoding *bla*_CTX-M-15_, among 7 other antimicrobial resistance genes was detected in 8 isolates belonging to 4 separate *K. pnuemoniae* STs: ST416, ST321, ST280, ST628, and *Enterobacter cloacae*, during a 1-year surveillance study analyzing rectal swab samples obtained from children in a paediatric oncology department in the Czech Republic [13]. Separately, the horizontal transfer of an epidemic IncFII/IncR *bla*_CTX-M-15_-containing plasmid, pK012_2 was detected in 48 different *K. pneumoniae* STs from stool samples collected from healthy and hospitalized children in the Dar es Salaam region in Tanzania [14]. The wide dissemination of a single plasmid across a series of unrelated bacterial clones implies ESBL-encoding plasmids bearing an F-like conjugation module constitute a well-adapted, low-fitness cost plasmid which can actively facilitate horizontal plasmid transfer, necessitating active surveillance.

In 2018, a clonal outbreak of *K. pneumoniae* ST628 was recorded in a UK district general hospital. The outbreak strain harbored a multi-drug resistant (MDR) plasmid, pESBL-PH encoding 11 AMR genes: *aac(3)-IIe*, *aac(6*′*)-Ib-cr*, *aph(3”)-Ib*, *aph(6)-Id*, *bla*_CTX-M-15_, *bla*_OXA-1_, *bla*_TEM-1B_, *dfrA14*, *qnrB1*, *sul2*, and *tet(A)*. Antibiotic susceptibility testing (AST) confirmed this plasmid was responsible for resistance to 6 drug classes including aminoglycosides, fluoroquinolones, cephalosporins, diaminopyrimidines, sulfonamides and tetracyclines [15]. Here, we sought to determine whether this plasmid existed in other *K. pneumoniae* STs, identifying whether similar plasmids to pESBL-PH-2018, the first plasmid identified in 2018 (March 3, 2018), isolated from a patient with a urinary tract infection (UTI), are present in the complete plasmid database, downloaded from NCBI, using strict thresholds.

## Method

### Genome Assembly and polishing

Bacterial strain UHD-2018 harboring pESBL-PH-2018 was assembled using hybrid read sets comprising illumina short reads and Oxford Nanopore Technology (ONT) long reads. Initially, raw illumina reads underwent quality control (QC) using fastp v.0.23.4 [16]. Here, low quality reads were discarded, and adapters and low-quality bases were trimmed using default settings. For long-read QC, a minimum length threshold of 6000-bp was applied to the ONT read set. NanoStat v.1.6.0 [17] was used to assess the N50 of the filtered read-set which yielded an N50 of 18,176-bp. Filtlong v.0.2.1 QC was again applied to the filtered read set to remove the worst 10% of the reads using the – keep_percent flag set to 90. Hybrid genome assembly was performed using Trycycler v.0.5.5 [18] using the filtered ONT long read sets using three different assemblers: Flye v.2.9.3-b1797 [19], miniasm/Minipolish v.0.1.2 [20] and Raven v.1.8.3 [21]. A consensus for each cluster generated was performed using trycycler consensus.

The ONT Medaka polishing tool v.1.11.3 was used to fix any potential remaining errors in the Trycycler ONT long-read assembly. The medaka model, r941_min_hac_g507 was used as it best matched the Nanopore pore and basecaller used for raw FAST5 basecalling. The Trycycler+Medaka assembly was subsequently polished using illumina short reads to fix small-scale errors (single-bp substitutions and indels) using Polypolish v.0.6.0 [22], yielding a FASTA file of the Trycycler+Medaka+Polypolish assembly for strain UHD-2018.

### Plasmid NCBI RefSeq database extraction

The outbreak plasmid, pESBL-PH-2018 was compared against the NCBI RefSeq Plasmid Database using the MinHash Algorithm for fast genome and metagenome distance estimation (Mash) [23] to enable efficient comparisons of large datasets. The complete NCBI RefSeq Plasmid Database, comprising 93,823 plasmids, was downloaded on July 16, 2024.

Mash similarity was determined as 1 – mash distance. Pairs of plasmids with mash similarity ≥ 0.98 have been considered the same plasmid previously [11]. Here, we applied this threshold, in addition to the number of shared hashes, to return similar completely assembled plasmids from the complete NCBI RefSeq database. Plasmids were filtered for both a shared hash ratio of 0.9 and a mash similarity ≥ 0.98, yielding plasmids with >91% coverage and >99.8% identity against pESBL-PH-2018. Using this strict threshold, highly similar plasmids can be captured, and we used this definition to define a pESBL-PH-2018-like plasmid. The accession number for each plasmid was used to retrieve the following information from NCBI: plasmid length, geographical source isolation, host strain, host disease, and host source.

### Plasmid antimicrobial resistance (AMR) genes, stress-related genes and in-silico conjugation detection

AMR, virulence and stress-associated genes were annotated using the AMRFinderPlus 2024-05-02.2 database [24]. Whole genome annotation of the assembled strain UHD-2018, including pESBL-PH-2018 was performed using Bakta v.1.9.3 [25] using the full database version, updated on July 16, 2024. In-silico plasmid conjugative capacity was predicted using Plascard [26] via the presence of a mobilization of plasmids (F) (MOBF) region, a type IV coupling protein (T4CP), and a type IV secretion system (T4SS).

### Single Nucleotide polymorphisms (SNPs) detection among recombination-filtered pESBL-PH-2018-like plasmids

Snippy v.4.6.0 (https://github.com/tseemann/snippy) was used to determine single-nucleotide polymorphisms (SNPs) between the annotated pESBL-PH-2018 plasmid and similar plasmids retrieved via the Mash search criteria described above. The reference Genbank file pESBL-PH-2018.gbff, output via the Bakta annotation tool, was set at the reference sequence. To determine the number of SNPs between the core genetic loci of pESBL-PH-2018 and other plasmids, predicted regions of recombination in other plasmids were removed using Gubbins v.3.3.1 [27]. The output file, recombination_predictions.gff was used to determine recombination regions.

Additionally, a strict threshold was applied to determine the relatedness between pESBL-PH-2018, and the query plasmids. An SNP count of fewer than 15 per 100-kbp was used to determine high similarity between the plasmids and infer likely horizontal plasmid transfer, as determined previously [28]. To achieve this, recombination regions were removed from the total query plasmid length. Core SNPs were then divided by the query plasmid length without the recombinant region(s). This returned a rate of SNPs per recombination free plasmid sequence, which was multiplied by 100,000 to give an SNP rate per 100,000.

### Plasmid detection in phylogenetically distinct bacterial clones

The host bacterial strain from each plasmid was downloaded from NCBI. A recombination free phylogenetic tree was constructed to confirm that strains harboring a pESBL-PH-2018 plasmid belong to phylogenetically distinct clones. Snippy-core v.4.6.0 (https://github.com/tseemann/snippy) was used to create an alignment between all host chromosomal FASTA files. Gubbins v.3.3.1 [27] was then run to create a recombination filtered tree using RAxML, generating a maximum-likelihood (ML) phylogenetic tree using 100 rounds of bootstrapping replicates. FigTree v1.4.4 was used to visualize the resulting phylogenetic tree.

### Core backbone evaluation

The core genome from plasmids derived from phylogenetically diverse host strains were evaluated to determine the core backbone and shared gene content. Roary v.3.12.0 [29] was used with the following parameters:

`roary -e –maftt, -p 8 *.gff`

## Results

### Plasmid Similarity against pESBL-PH-2018: an overview

The strict search criteria yielded 61 similar plasmids. The similarity between the query plasmids and the reference pESBL-PH-2018 was compared using a series of methods. The Mash distance metric, based on the Jaccard index of *k*-mer sketches (subsequences of length *k*) was initially used to assess plasmid similarity (Supplementary File 1). A mean Mash similarity (1 – mash distance) of 0.9992; standard deviation (σ): 0.0007, IQR: 0.9988-0.999 was recorded. Additionally, the number of shared hashes was assessed as a ratio. Across the collection of 61 plasmids samples, the hash ratio ranged from 0.902-1.00 (mean: 0.968, IQR: 0.954-0.998, median: 0.969, σ: 0.029).

The nucleotide coverage and identity were also compared between each of the 61 query plasmids and the reference, pESBL-PH-2018. BLASTn coverage for the plasmids against pESBL-PH-2018 ranged from 91-100% (mean: 97.82%, IQR: 97-100%, median: 98.0%, σ: 2.34). The BLASTn identity of the query plasmids against pESBL-PH-2018 was high; range 99.87-100.00% (mean: 99.99%, IQR: 99.99-100.00%, σ: 0.019). Finally, the length of the plasmids was assessed. The 61 query plasmids had a mean length of 244,365-bp. When each plasmid was compared against the reference length of pESBL-PH-2018, 247,179-bp, the IQR for plasmid length was between 97.56-99.99% of the length of pESBL-PH-2018. Taken together, the high Mash similarity, shared hashes, high coverage and nucleotide identity, combined with a similar length, suggests all 61 plasmids are highly similar to pESBL-PH-2018.

### Identification of pESBL-PH-2018-like plasmids: Host strain and isolation source

The host bacterium strain harboring each of the query plasmids were downloaded from the NCBI nucleotide database. Across the collection of 57 plasmid samples where a host bacterium ST designation was assigned, 13 unique sequence types (STs) were identified. The pESBL-PH-2018-like plasmid was detected most frequently in *K. pneumoniae* ST307 (*n*=21/57, 36.84%). Furthermore, pESBL-PH-2018-like plasmids were found 5 STs where the ST was identified 3 or more times: ST628 (*n*=9/57, 15.78%), ST323 (*n*=8/57, 14.03%), ST29 (*n*=5/57, 8.77%), ST405 (*n*=3/57, 5.26%), and

ST11 (*n*=3/57, 5.26%). In addition, the pESBL-PH-2018-like plasmid was identified in 7 other STs: ST617, -280, -479, -6265, -2856, and ST584, respectively (Figure 1A). Across 60 samples where a host strain species was recorded, the pESBL-PH-2018 like plasmid was predominately detected in *K. pneumoniae* (*n*=57/60, 95%). Separately, three similar plasmids were detected in two different genera, while a single plasmid was detected in a different species; plasmid P1 from a *Klebsiella oxytoca* strain, plasmid 2 from KPN029 *Klebsiella variicola* ST347, while an *Escherichia coli* ST479 strain carried pCUVET16-321.1. Together, these results suggest different *Klebsiella* genera and other species may be receptive carriers for pESBL-PH-2018-like plasmids.

**Figure 1.**
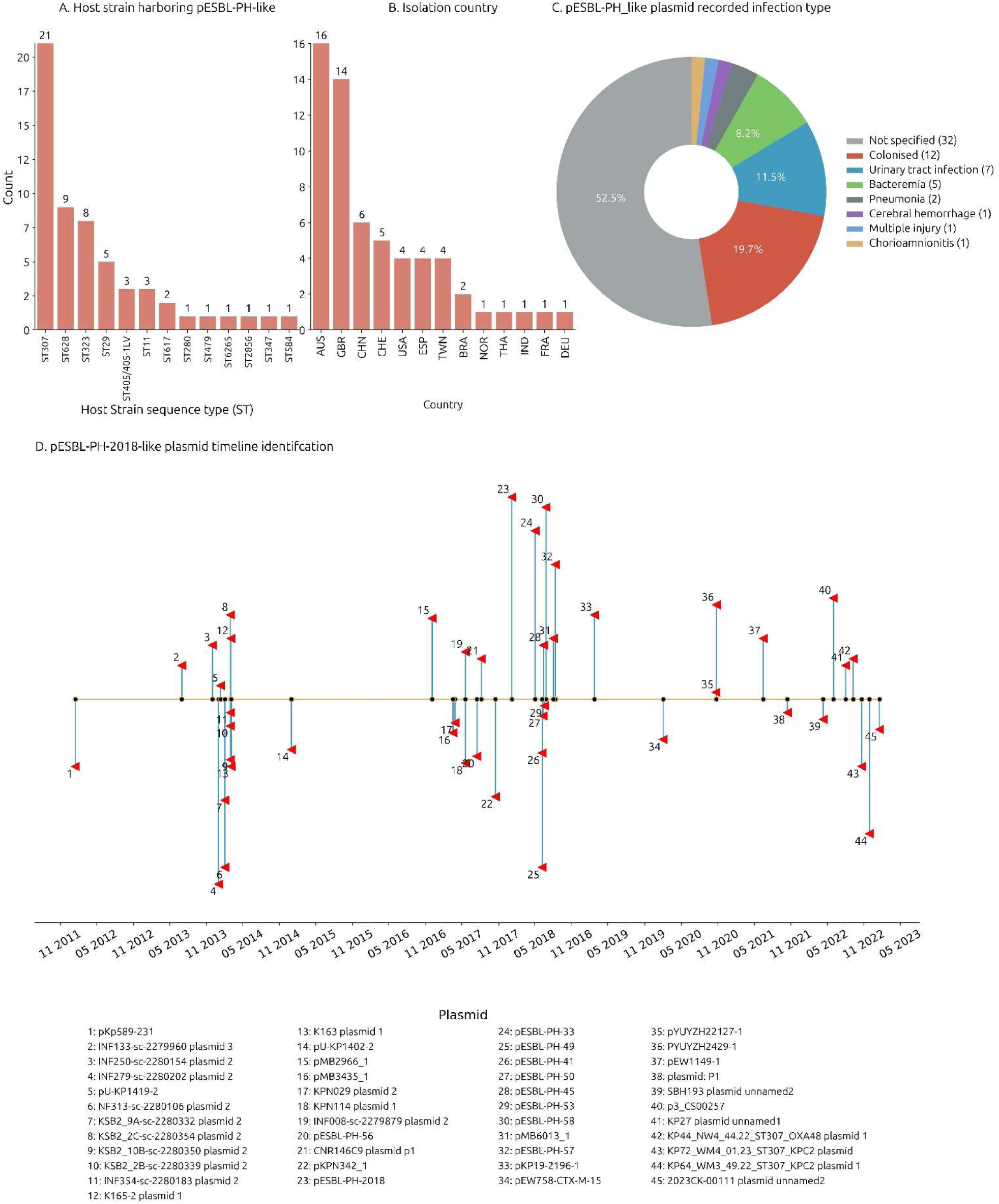
Plasmid host strain, country, infection source and timeline identification. A. Host strain ST designation. B. Country of isolation accessed via available Biosample data from NCBI. C. Infection type assignment determined according to meta-data available via Biosample data from NCBI. D. The identification of similar plasmids to pESBL-PH-2018 plotted as per their discovery. Each plasmid name matches the numbered key. Plasmid meta-data is available in Supplementary file 1.

The pESBL-PH-2018-like plasmid was identified in 13 unique countries between 2012-2023 (Figure 1B). Available Biosample data from NCBI indicated the plasmid (including pESBL-PH-2018) was most often found in Australia (26.22%, *n*=16/60), the United Kingdom (22.95%, *n*=14/60), and China (9.83%, *n*=6/60). Separately, the plasmid was identified ≥ 4 times in Switzerland, USA, Spain, and Taiwan (Figure 1B). Available Biosample data also indicated the pESBL-PH-2018-like plasmids (including pESBL-PH-2018) were identified from a range of infection types including urinary tract infections (11.3%, *n*=7), bacteremia (8.1%, *n*=5), pneumonia (3.2%, *n*=2) cerebral hemorrhage (1.6%, *n*=1), acquisition from wounds via injuries (1.6%, *n*=1), and chorioamnionitis (1.6%, *n*=1), respectively (Figure 1C).

The collection dates of plasmids with a recorded year, month and day were investigated. Plasmids similar to pESBL-PH-2018 were collected from January 16, 2012, from Barcelona, Spain (pKp589-231, accession: CP028817.1) through to January 17, 2023, from the USA (2023CK-0011, plasmid unnamed, accession: CP131947.1). The interval between the detection of these two plasmids is 11 years and 4 days. Similar plasmids were detected from 25% (*n*=11) of the samples with a collection date in 2014, while 20.45% (*n*=9) were collected in 2018, however ≥ 3 similar plasmids were recorded in 2014, 2017, 2018, 2020, and 2022 (Figure 1D). These results reveal a consistent presence of ESBL-encoding plasmids similar to pESBL-PH-2018. A recombination-filtered phylogenetic tree of host bacterial strains revealed the plasmids were carried by distinct bacterial clones. Across these clones, evidence of expansion of the host strain with the plasmid is seen (Figure 2).

**Figure 2.**
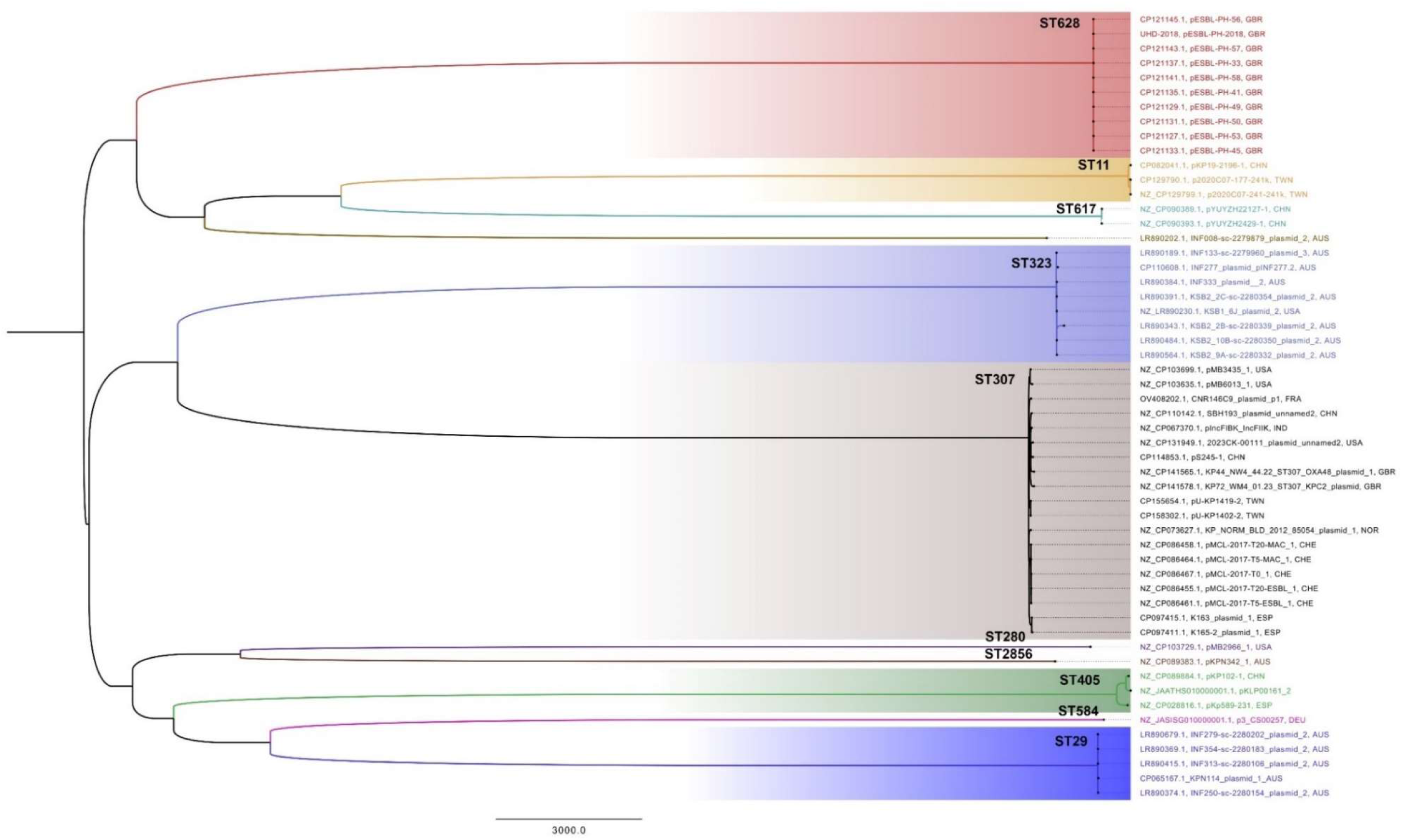
Host strain carrying plasmids similar to pESBL-PH-2018 colored according to their clade. Plasmid accession, name, and country source are listed at the tips of each branch. The recombination-filtered phylogenetic tree was constructed using RAxML, generating a maximum-likelihood (ML) phylogenetic tree using 100 rounds of bootstrapping replicates. FigTree v1.4.4 was used to visualize the resulting phylogenetic tree. The scale refers to the number of SNPs.

### Single nucleotide polymorphism analysis

Single nucleotide polymorphisms (SNPs) between pESBL-PH-2018 and the query plasmids were investigated to provide high-resolution analysis of their relatedness. SNP analysis was performed on recombinant-removed query plasmids. Across the selection of plasmids, an average of 4.95 SNPs were detected (range: 0-16, IQR: 3-6). 93.44% (*n*=57/61) plasmids had an SNP count ≤ 10, while 65.5% (*n*=40/61) had a SNP count of ≤ 5. With the exception of CNR146C9 plasmid 1 and p3_CS00257 which each had a recombination region of 6,757-bp, and 17,782-bp respectively, a mean recombinant region of 531-bp was identified (range: 0-2,396-bp), which only constituted on average 0.22% of their overall plasmid length, revealing the plasmids share large stretches of identical sequences. Across all plasmid samples, an average of 1.95 SNPs/100-kb was identified, suggesting the plasmids are closely related to pESBL-PH-2018 (Figure 3).

**Figure 3.**
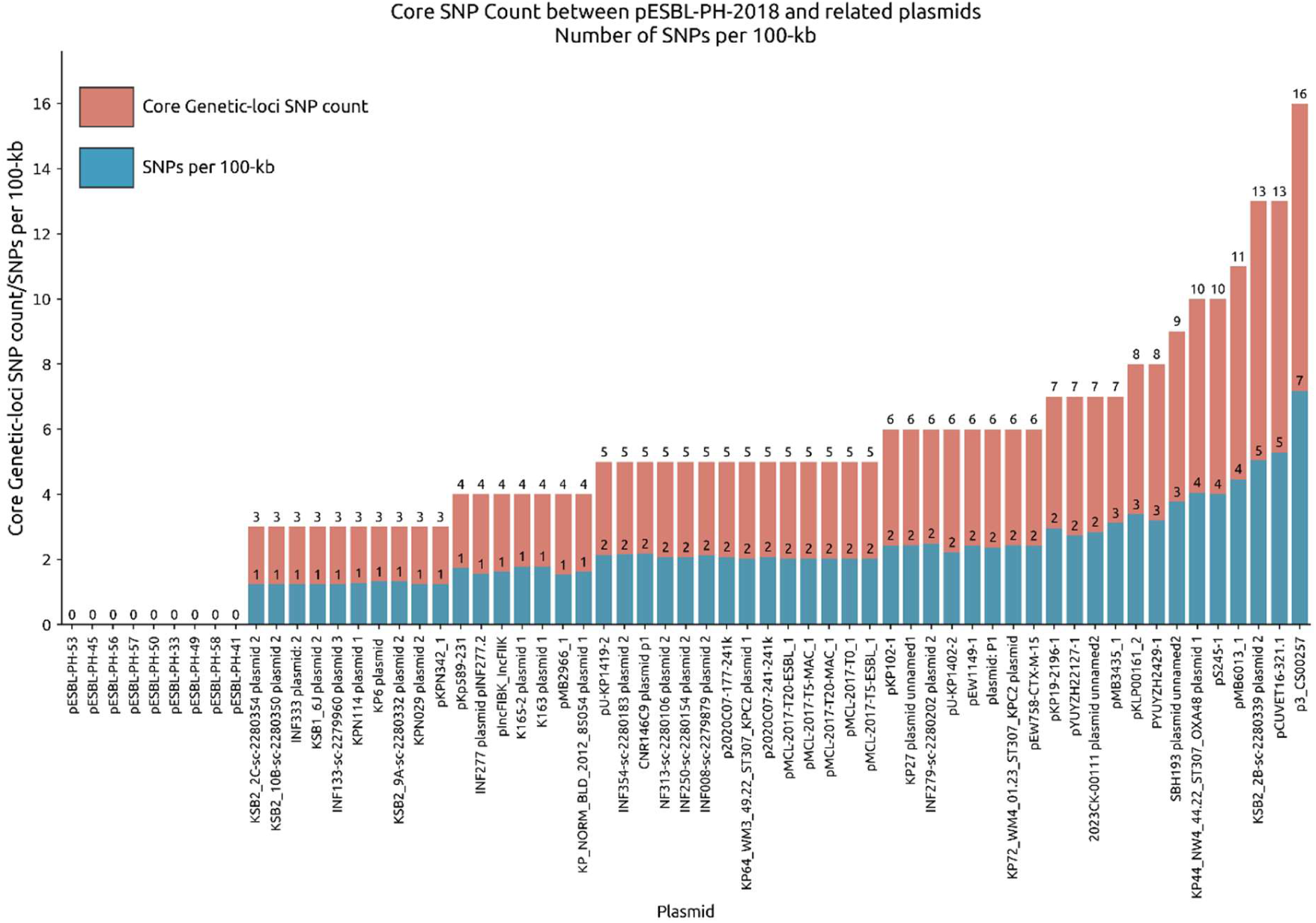
SNP distance between pESBL-PH-2018 and query plasmids. The core genetic loci between query plasmids and pESBL-PH-2018 were compared to determine the number of SNPs to use as a proxy of plasmid relatedness (red). Plasmid relatedness was normalized to the number of SNPs/100kb in core genomic loci (blue).

### Core genome analysis

The core genome for 12 plasmids (Table 1), identified in 12 unique STs was evaluated to determine the core backbone and shared gene content between plasmids from diverse host strains. Across the 12 plasmids, 210 genes were shared between all plasmids. The total nucleotide length from these 210 shared genes was 182,216-bp. In addition, 90.9% of genes (*n*=281/309) were present in either 99-100% (core genes), or 15-95% (shell genes) from the 12 plasmids, while only 9.1% (28/309) cloud or accessory genes were determined. A plasmid backbone for a selection of representative STs is shown in Figure 4. While the plasmid analysis yielded 13 unique host strain STs, *K. pneumoniae* strain ST280 carrying pMB2966_1 was excluded from the core genome analysis due to the presence of a segmental duplication harboring *sul2*, *aph(3’’)-Ib*, *aph(6)-Id*, *bla*_TEM-1_ and *bla*_CTX-M-15_, respectively. All plasmids shared several stress-related genes; 100% identity and coverage were identified for silver (*silS*, *silR*, *silC*), copper (*pcoABCR*), and arsenic (*arsABCR*) heavy metal resistance genes, alongside the heat shock gene, *hsp20*, while all plasmids were typed as IncFIB(K). All plasmids are predicted to be conjugative, possessing components necessary for conjugation, including a relaxase, T4SS, and a T4CP.

**Figure 4.**
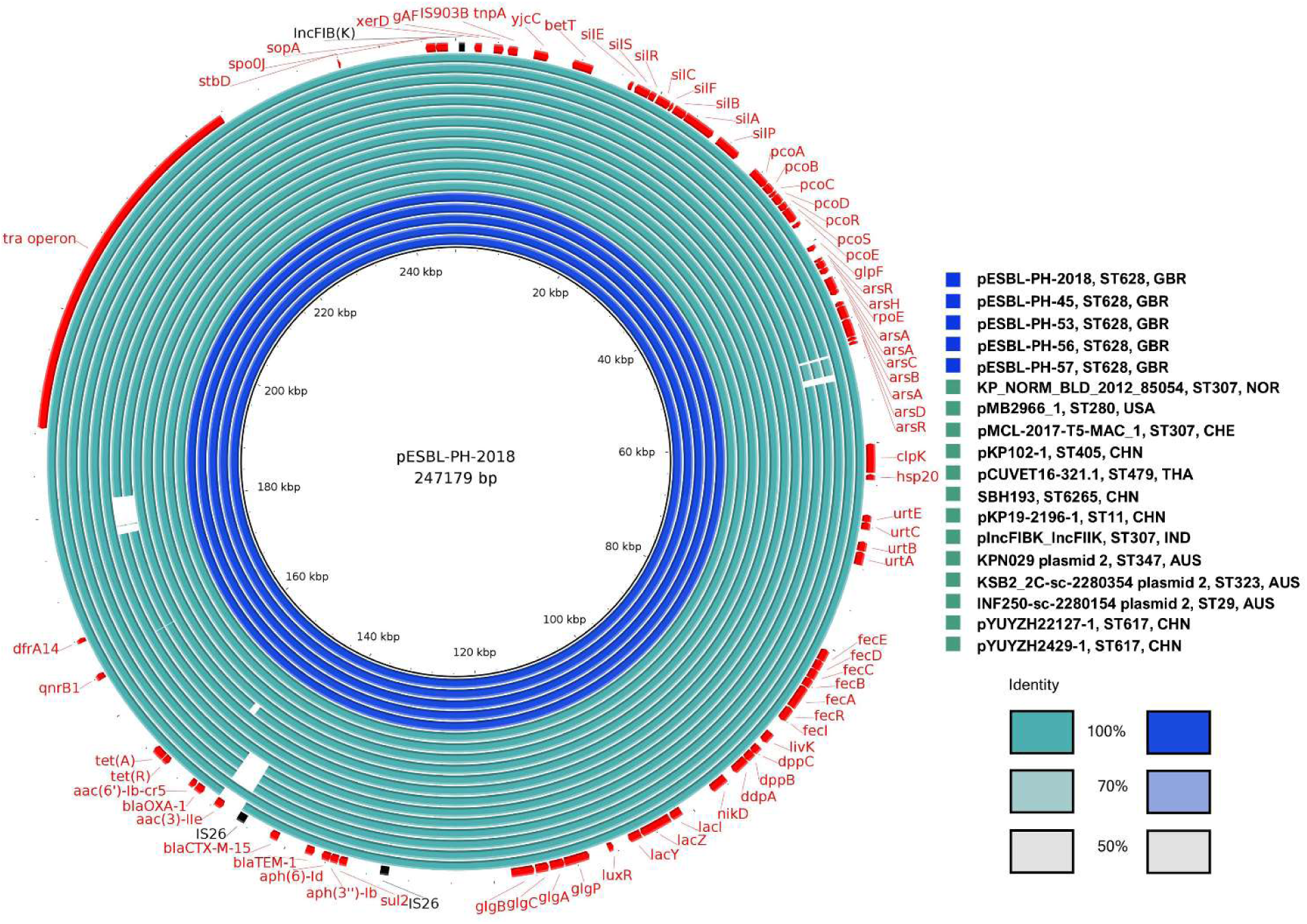
Comparison of pESBL-PH-2018 with similar plasmids. Each plasmid is represented by a circular ring, ordered from the innermost (top of the key), to the outermost (bottom of the key). The plasmid visual comparison was performed using the BLAST Ring Image Generator (BRIG) [30]. Comparison was performed across a range of plasmids identified from 11 unique hosts. Gene are indicated on the outer ring in red with their direction of transcription indicated by an arrowhead.

**Table 1.** Plasmid samples used for core genome analysis.

| Plasmid | Host ST | Host Species |
| --- | --- | --- |
| KPN029 plasmid 2 | ST347 | *K. variicola* |
| KP_NORM_BLD_2012_85054 plasmid 1 | ST307 | *K. pneumoniae* |
| pKP19-2196-1 | ST11 | *K. pneumoniae* |
| pKPN342_1 | ST2856 | *K. pneumoniae* |
| pKP102-1 | ST405 | *K. pneumoniae* |
| pYUYZH22127-1 | ST617 | *K. pneumoniae* |
| SBH193 plasmid 2 | ST6265 | *K. pneumoniae* |
| pESBL-PH-56 | ST628 | *K. pneumoniae* |
| pCUVET16-321.1 | ST479 | *E. coli* |
| P3_CS00257 | ST584 | *K. pneumoniae* |
| INF250-sc-2280154 plasmid 2 | ST29 | *K. pneumoniae* |
| KSB2_2C-sc-2280354 plasmid 2 | ST323 | *K. pneumoniae* |

### AMR variants

From the collection of plasmids including pESBL-PH-2018, 54.8% (*n*=34/62) carried exactly one copy of the 11 AMR genes: *aph(3”)-Ib*, *aph(6)-Id*, *bla*_TEM-1B_, *bla*_CTX-M-15_, *aac(3)-IIe*, *bla*_OXA-1_, *aac(6*′*)-Ib-cr*, *tet(A)*, *qnrB1*, *dfrA14* and *sul2*. Plasmids pMB2966_1 and pMB6013_1 carried multiple copies of these genes, including either 2 or 3 copies each of the ESBL gene, *bla*_CTX-M-15_, respectively (Figure 5). In total, 58% of the plasmids carried either 1 or more of the total complement of 11 AMR genes carried by pESBL-PH-2018. Notably, a common variant, which lacked only *aac(3)-IIe* was detected in 24.19% (*n*=15/62) of samples. From these 15 samples, 13 samples were identified from Australia, confirming, the *aac(3)-IIe* aminoglycoside gene was commonly missing from plasmids related to pESBL-PH-2018 sourced from Australia.

**Figure 5.**
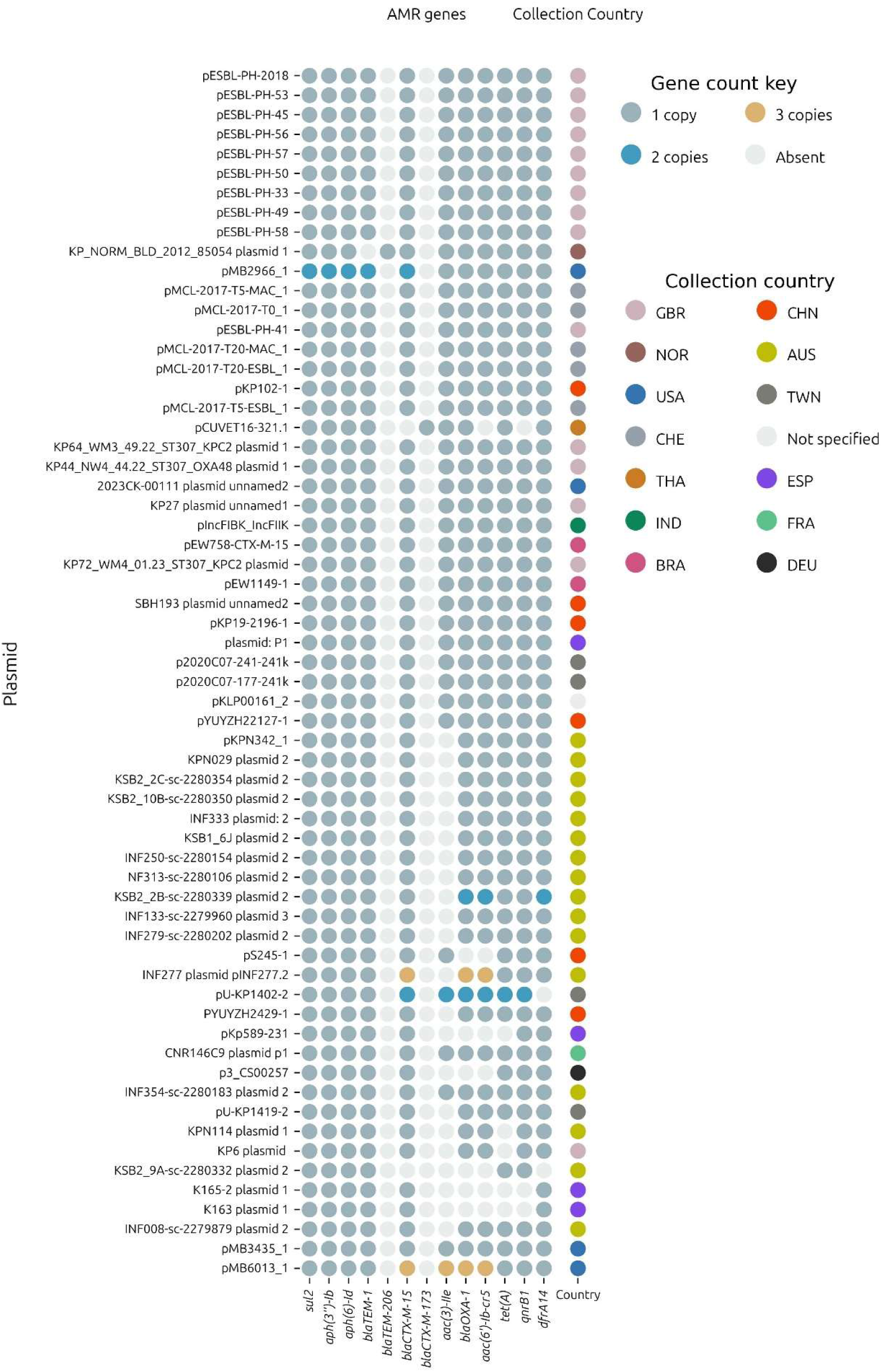
Antimicrobial resistance genes present among plasmids similar to pESBL-PH-2018. AMR genes are marked as absent (white), and according to their copy number: 1 (grey), 2 (blue), and three (light brown). Countries are indicated according to their IBAN country code and colored.

Although similar plasmids to pESBL-PH-2018 were detected, a number of variants with a different number of the same AMR genes were observed. The aminoglycoside resistance gene, *aac(3)-IIe*, was absent from a number of plasmids. In comparison to pESBL-PH-2018, plasmid pKPN342_1 (accession: CP089384.1), an AMR variant lacking *aac(3)-IIe*, had a deletion of 3,420-bp including *aac(3)-IIe* and a complete copy of IS*26* located between 2 inverted sequences (Figure 6). A similar deletion of 2,600-bp encompassing *aac(3)-IIe* was identified in plasmid KBS2_10B (accession: LR8900485.1), however in this plasmid both copies of the IS*26* between *aac(3)-IIe* were retained.

**Figure 6.**
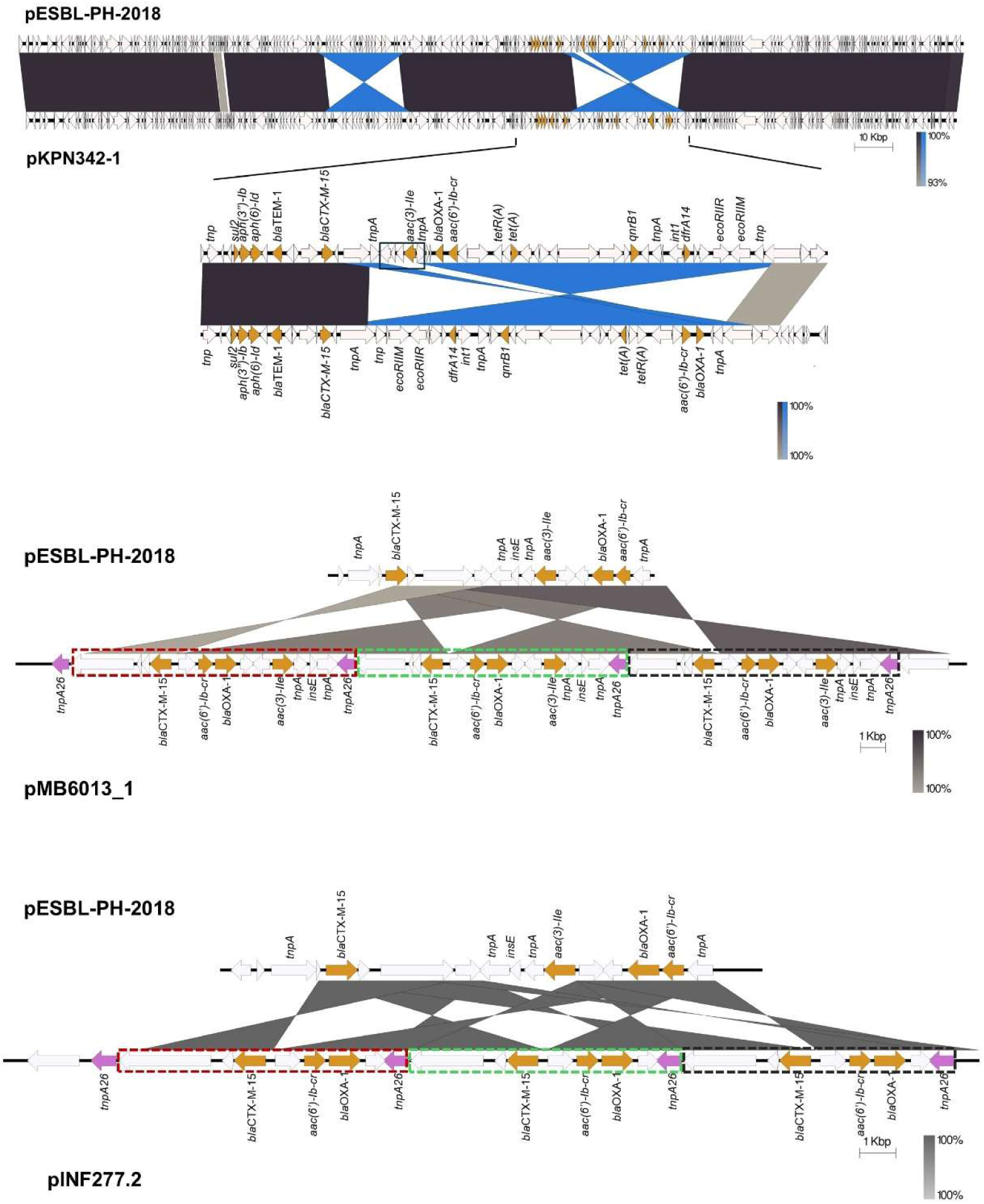
**Comparison between the AMR region of pESBL-PH-2018 and similar plasmids**. Top. Likely IS*26*-medatied deletion of the aminoglycoside resistance gene, *aac(3)-IIe*. Middle. For pMB6013_1 (accession: CP103636.1) isolated from a *K. pneumoniae* ST307 strain from a patient with bacteremia, three tandem copies of the AMR genes: *bla*_CTX-M-15_, *aac(6’)-Ib-cr*, *bla*_OXA-1_, *aac(3)-IIe* were present in a 11,009-bp array. For pINF277.2, IS*26*-mediated TU amplification of a 7,559-bp region carrying the AMR genes: *bla*_CTX-M-15_, *aac(6’)-Ib-cr*, *bla*_OXA-1_, *aac(3)-IIe* was present in a tandem array as indicated by the three colored bordered boxes. Tandem repeats were measured from the inverted repeat left (IRL) of IS*26* to the end of the shared Tn3 family transposase fragment, represented in each tandem array as the longest white arrow (at the left end of each bordered box).

Separately, KP_NORM_BLD_2012_85054 plasmid 1 (accession: CP073628.1) from *K. pneumoniae* ST307, encoded the TEM variant, *bla*_TEM-206_. This variant was due to an SNP at nucleotide position 25, resulting in a missense variant, whereby the non-polar amino acid alanine has been replaced by the polar amino acid threonine at amino acid position 9.

Variants with 2 copies of a number of AMR genes, including the ESBL gene, *bla*_CTX-M-15_ were identified. pMB2966_1 had 2 copies each of the following genes: *sul2*, *aph(3’’)-Ib*, *aph(6)-Id*, *bla*_TEM-1_ and *bla*_CTX-M-15_, respectively. In pESBL-PH-2018, these AMR genes are present within a 15,310-bp IS*26* PCT. However, in pMB2966_1 a segmental duplication, (mobilization to another genomic context), of a 16,374-bp region, including an inverted copy, carrying these genes was observed. Both the regions carrying these AMR genes shared 12,212-bp with the IS*26* from pESBL-PH-2018, but both segments lacked IS*26* insertions sequences bracketing the 5 AMR genes, confirming the duplication of these genes did not occur due to an IS*26*-mediated translocatable unit (TU) amplification. Additionally, plasmid 2 (accession: LR890344.1) from *K. pneumoniae* ST323 obtained from a rectal swab had a segmental duplication of 10.98-kb, including the AMR gene *dfrA14*. Similar to pMB2966_1, the second segmental copy lacked IS*26* elements either side of the duplication.

For plasmid pU-KP1402-2, 2 copies each of *bla*_CTX-M-15_, *aac(3)-IIe*, *bla*_OXA-1_, *aac(6’)-Ib-cr*, *tet(A)*, and *qnrB1* were identified. Here, *bla*_CTX-M-15_ and *aac(3)-IIe* were duplicated as an additional inverted copy associated with a single copy of IS*26 tnpA26*. Another inverted duplication was also observed for *bla*_OXA-1_ and *aac(6’)-Ib-cr* with both copies having inwardly facing *tnpA26* transposases. Interestingly, both *tet(A)* and *qnrB1* were also duplicated.

Two plasmid variants, pMB6013_1 (accession: CP103636.1), and pINF277.2 (accession: CP110610.1) with a copy number of three for the ESBL gene *bla*_CTX-M-15_ and other AMR genes were also investigated. Here, IS*26*-TU tandem amplification was identified. While pESBL-PH-2018 was similar to pMB6013_1, the AMR region varied. Relative to pESBL-PH-2018, a transposase gene and the ESBL gene *bla*_CTX-M-15_ are inverted. This inversion lies upstream of an IS*26 tnpA26* and downstream of an inverted region carrying three AMR genes, *aac(6’)-Ib-cr*, *bla*_OXA-1_, and *aac(3)-IIe*. Together, this forms a section of 4 AMR genes bracketed by IS*26 tnpA26 on* each side (Figure 4).

This region has duplicated, generating three tandem copies of these AMR genes. The AMR tandem repeat is 11,009-bp, measured from the inverted repeat left (IRL) of IS*26* to the shared region of the Tn3 transposase fragment (marked boxes Figure 6). The genomic organization indicates IS*26*-mediated TU amplification has created the tandem array.

Separately, pINF277.2 from *K. pneumoniae* ST323, isolated from a bronchoalveolar lavage (BAL) sample from a patient with pneumonia also had a similar IS*26*-medaited TU amplification for the three AMR genes: *bla*_CTX-M-15_, *aac(6’)-Ib-cr*, and *bla*_OXA-1_. Relative to pESBL-PH-2018, *bla*_CTX-M-15_ is inverted and lies upstream of a shared IS*26 tnpA26* transposase. The AMR region from pESBL-PH-2018 including *aac(6’)-Ib-cr*, *bla*_OXA-1_ and *tnpA26* is inverted and located downstream of *bla*_CTX-M-15_ (Figure 6). This region in pINF277.2, bracketed by *tnpA26* is duplicated three times, increasing the copy number for each of these genes to three. The IS*26*-mediated TU tandem array here was 7,559-bp (marked boxes Figure 6).

### Host strain background

The host strain background for 56 strains with a complete chromosome was investigated for AMR, virulence and stress-associated factors. In particular, gene mutations associated with carbapenem resistance were analyzed. The pESBL-PH-2018-like plasmid was present in a background associated with carbapenem resistance mechanisms. Notably, 26.78% (*n*=15/56) of the strains encoded a carbapenem resistance gene on an accessory plasmid in addition to carrying a pESBL-PH-2018-like plasmid (Figure 7). In addition, 17.85% (*n*=10/56) of the strains had a truncation in the outer membrane porin gene, *ompK36*, disruptions of which can reduce the permeability of the outer membrane to carbapenems. Porin truncations were present in 4 strains which also carried a carbapenemase gene on an accessory plasmid (Figure 7). Three strains carried the *ompK36* variant, *ompK36_D135DGD*, associated with porin constriction mediated by the di-amino acid (Gly115-Asp116) insertion into loop 3 of the OmpK36 porin [31]. This change has been associated with restricted access of carbapenems into the host cell. Two of these variants were found in addition to carbapenem resistance genes presents on separate plasmids (Figure 7, Supplementary file 2).

**Figure 7.**
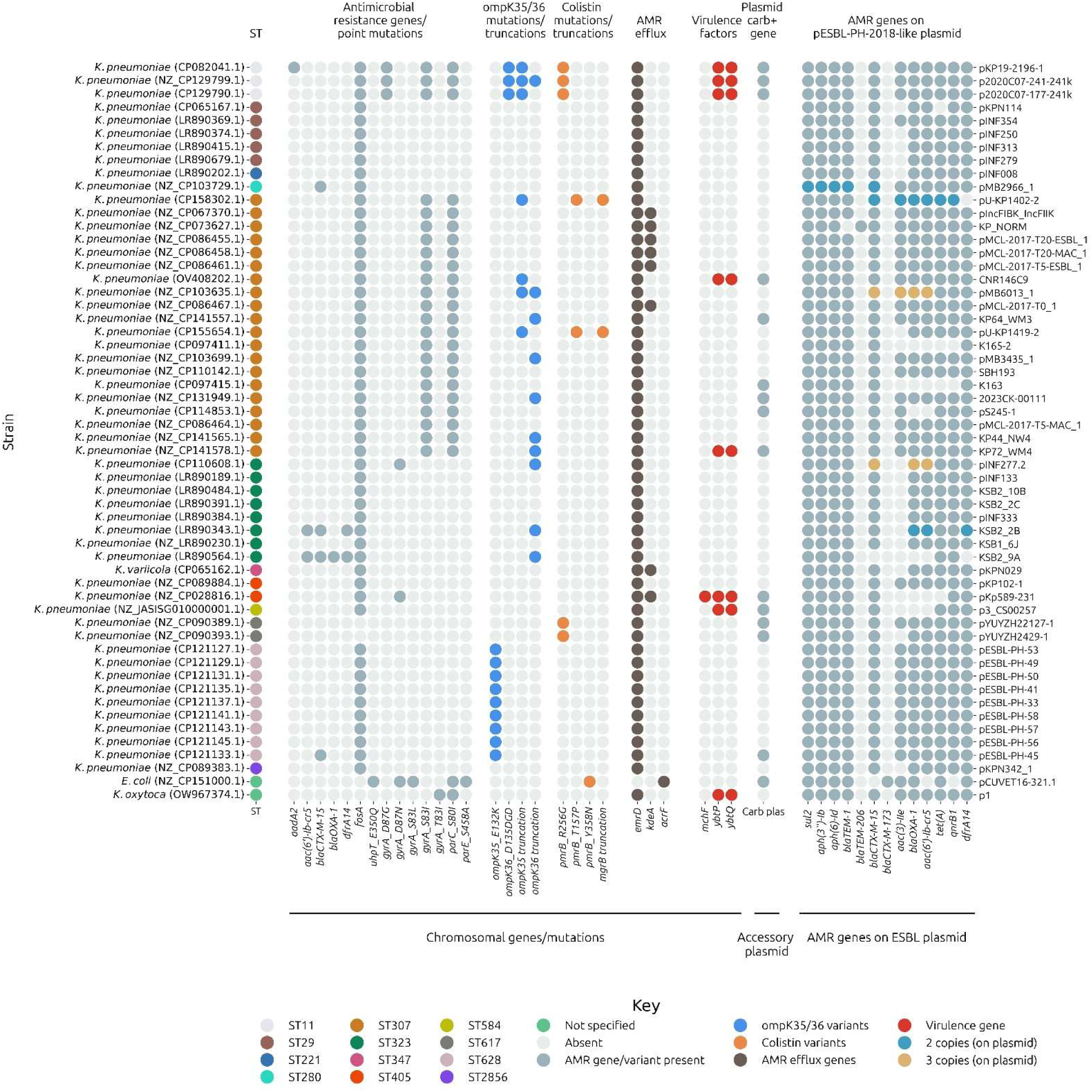
AMR genes/point mutations, porins and virulence associated factors among strains carrying pESBL-PH-2018-like plasmids. Strains carrying a pESBL-PH-2018-like plasmid are listed on the left y-axis according to their ST; the pESBL-PH-2018-like plasmid carried by the strain is indicated on the right y-axis. In the AMR section, *aadA2*, and *aac(6’)-Ib-cr* confer resistance to aminoglycosides, while *bla*_CTX-M-15_ confers resistance to beta-lactam antibiotics, particularly third-generation cephalosporins, *bla*_OXA-1_ provides resistance against beta-lactam antibiotics including penicillins and some narrow-spectrum cephalosporins. The *dfrA14* confers resistance to trimethoprim, while the *fosA* gene, and the *uhpT_E35Q* gene variant confer resistance to fosfomycin. *gyrA* and *parC* gene variants are associated with resistance to fluoroquinolones. *ompK35/36* porin variants are associated with a rise in the MIC for carbapenems, while *pmrB/mgrB* gene variants provide resistance against colistin. The chromosomal genes/mutations bar at the bottom refers to AMR determinants on the chromosome, the accessory plasmid bar refers to plasmids separate to pESBL-PH-2018-like carrying a carbapenemase gene, while the AMR genes on the ESBL plasmid bar refer to resistance genes located on plasmids similar to pESBL-PH-2018.

Colistin resistance associated gene variants were also detected in 14.28% (*n*=8/56) strains. Here, 5 strains carried the *pmrB* gene variant: *pmrB_R256G*, 1 carried the *pmrB_Y358N* variant, while 2 strains carried both the *pmrB_T157P* alongside a gene truncation in *mgrB*. Such changes have been associated with constitutive activation of the two-component system, PmrA-PmrB, leading to a reduction in the negative charge of the bacterial lipopolysaccharide (LPS) present on the bacterial outer membrane. The reduction in negative charge decreases the binding affinity of colistin (a cationic peptide), leading to colistin resistance.

Notably, a large proportion; 92.85% (*n*=52/56) of the isolates were predicted to encode a chromosomal fosfomycin resistance gene, *fosA*. In addition, a number of predicted quinoline resistance inducing mutations impacting either DNA gyrase, or topoisomerase IV were detected. In particular, all representative ST11, and ST307 isolates encoded the *gyrA* S83I, and *parC* S80I double mutations. A proportion of strains, 14.28% (*n*=8/56) also encoded the virulence-associated YbtPQ ABC transporter, a key transporter system involved in both iron acquisition during infection and broad export of antimicrobials. Taken together, AMR, virulence and stress associated typing of the chromosomes and plasmids, co-present with the pESBL-PH-2018-like plasmid reveal a genomic background associated with genes conferring resistance to critical and last-resort antimicrobials such as carbapenems and colistin, in addition to the 6 classes of AMR resistance conferred by pESBL-PH-2018.

## Discussion

Multiple lines of evidence demonstrate the widespread dissemination of a pESBL-PH-2018-like plasmid. These include a high Mash similarity, high coverage and nucleotide identity, and a low SNP difference in the core genetic loci of query plasmids. Similar plasmids also carried either identical or a similar number of the same AMR genes, alongside an identical plasmid replicon, heavy metal resistance genes and conjugative transfer genes. The broad-host range of CTX-M-15 encoding plasmids has been recorded previously [11,14]. In addition, expansion of different bacterial clones harboring a similar CTX-M-15 encoding plasmid has been reported [32,33]. pESBL-PH-2018 may represent another plasmid which can transfer between hosts. Once transferred, clonal expansion within receptive hosts can occur.

Plasmids similar to pESBL-PH-2018 were identified from 2012-2023, further confirming the long-term persistence of a pESBL-PH-2018-like plasmid. High conjugation efficiency has previously been reported for CTX-M-15 encoding plasmids, with high conjugation frequencies in the order of 10^-1^ recorded [34]. Together, the presence of similar plasmids in distinct hosts, alongside the identification of conjugative transfer genes may indicate horizontal gene transfer (HGT) may be responsible for the widespread dissemination of a pESBL-PH-2018-like plasmid. However, separate conjugation assays are required to determine the conjugative frequency for pESBL-PH-2018 and similar plasmids.

The AMR region varied between the plasmids. Notably, the aminoglycoside resistance gene *aac(3)-IIe* was absent on a number of plasmids and may have occurred via IS*26*-mediated deletion, as strains lacked both the resistance gene and IS*26* (such as, pKPN342-1). For two plasmids, pMB6013_1, and pINF277.2, the copy number of the ESBL gene, *bla*_CTX-M-15_ alongside other AMR genes was repeated three times in a tandem array. Antibiotic selection pressure can lead to the formation of TUs, which may incorporate into either the original plasmid location or a distinct location and amplify via rolling circle replication. Tobramycin selection pressure has been shown to lead to the duplication of the *aphA1* kanamycin and neomycin resistance gene from an *Acinetobacter baumannii* clinical strain MRSN56, due to IS*26*-mediated TU formation [35]. The AMR region of pESBL-PH-2018 may be subject to rearrangements and resistance gene duplications due to the presence of IS*26*, combined with antibiotic selection pressure often encountered in clinical settings. Concerningly, for both plasmids with tandem repeats of AMR genes, dual β-lactamase-encoding genes, *bla*_CTX-M-15_ and *bla*_OXA-1_ were identified. IS*26*-mediated co-amplification of two β-lactamase-encoding genes can yield a phenotype which is often non-susceptible to carbapenems such as meropenem or ertapenem [36].

Beyond the presence of AMR and virulence genes present on the pESBL-PH-2018-like plasmid, the genotypic background of these strains was investigated to identify additional resistant/virulence-linked determinants associated with carbapenem and colistin resistance. A proportion of the strains carried the chromosomal integrative and conjugative element ICE*Kp* encoding the ABC transporter YbtPQ. Recently, the YbtPQ transporter has been shown to reduce bacterial susceptibility to several classes of antibiotics, including carbapenems, likely mediated via broad range antimicrobial export [37]. The co-presence of carbapenemase genes in 7/8 of the strains encoding YbtPQ may increase the minimum inhibitory concentration (MIC) for carbapenems, and other classes of antibiotics while contributing towards enhanced virulence via iron acquisition.

Furthermore, disruptions in the outer membrane porin gene, *ompK36* in concert with the presence of β-lactamase genes have been linked with the development of non-carbapenemase producing carbapenem resistant Enterobacterales (non-CP-CRE). For example, a non-CP-CRE *K. pneumoniae* isolate, MB101, carried tandem amplification of a TU carrying *bla*_OXA-1_ together with an IS*Ecp1* transposition unit inserted into *ompK36* [38]. Six strains carried *ompK36* truncations without the presence of either chromosomal or plasmid carbapenemase genes. Notably, in two strains, multiple plasmid *bla*_CTX-M-15_ genes were detected with either *ompK36* truncations (strain 3221, chromosomal accession: NZ_CP103635.1), or *ompK35* (strain U-KP1402, chromosomal accession: CP158302.1), yielding a potential non-CPE genotype. These two strains, and the other 4 strains with a single *bla*_CTX-M-15_ copy, may have a background primed for carbapenem resistance development, a worrying scenario based on the presence of CTX determinants which already confer resistance to cephalosporins, and other strains which already carried carbapenemase genes on separate plasmids. Such strains will be harder to treat and may present more opportunities for onward ESBL plasmid dissemination.

Low levels of antibiotics or heavy metals in the polluted external environment or antibiotic consumption may promote the selection and enrichment of MDR plasmid carriage among bacteria. Concerningly, very low concentrations of single antibiotics or heavy metals, or combinations of these compounds promote the selection for a large MDR 220-kbp plasmid, pUUH239.2 that carries resistance towards aminoglycosides, β-lactams, tetracycline, macrolides, trimethoprim, sulfonamide, silver, copper, and arsenic [39]. Both antibiotic and heavy metals at sub-minimum inhibitory concentrations (MICs) allow selection of the MDR plasmid, pUUH239.2; indeed, the minimum selective concentration (MSC) for tetracycline was 17-fold lower, while the MSC for arsenite was approximately 140-fold lower than the MIC of the same susceptible bacterial strain lacking pUUH239.2 [39]. Areas impacted by human pharmaceutical use may promote MDR plasmid maintenance and enrichment and may be an important factor governing long-term persistence and dissemination of ESBL-encoding plasmids which similarly encode antimicrobial and heavy metal resistance determinants, such as the pESBL-PH-2018-like plasmids. For example, in European surface waters, trimethoprim has been detected in concentrations up to several hundred ng/L [40], exceeding the MSC for trimethoprim.

Using the strict thresholds set, we identified plasmids similar to pESBL-PH-2018 detected during a nosocomial outbreak in the United Kingdom. The strict thresholds applied however, may underestimate the true prevalence of the ESBL-encoding plasmid. For example, we recently identified an ESBL encoding plasmid, pEBM1 which underwent co-integrate formation with another *bla*_CTX-M-15_ encoding plasmid from a *K. pneumoniae* strain [41]. A plasmid co-integrate may not be recognized as highly similar using the similarity thresholds. Our approach was designed to detect very similar plasmids, with a similar number of AMR genes; only 4 plasmids detected were missing 4 or less AMR genes from the 11 present on pESBL-PH-2018. However, despite this, the strict thresholds applied, definitively prove plasmids very closely related to pESBL-PH-2018 have a wide host range, including across bacterial genera and species, and display remarkable conservation over a 10-year period, existing in diverse strains with an extensive drug-resistant profile.

## Conclusion

A plasmid identified from an outbreak strain of *K. pneumoniae* ST628, was structurally similar to plasmids identified in phylogenetically distinct *K. pneumoniae* hosts strains. The plasmid was also identified in different *Klebsiella* genera and *E.coli*. Similar plasmids across distinct strains suggests the plasmid may disseminate via horizontal plasmid transfer into recipient hosts. Expansion in these hosts indicates the plasmid imposes a limited fitness burden. Furthermore, the combination of AMR genes with IS*26* may provide an environment whereby the copy number of ESBL genes can amplify, as detected in two strains here. Finally, the ESBL-encoding plasmid was identified in a background of AMR and virulence determinants. Importantly, in addition to ESBL plasmid carriage, truncations in the porin *ompK36*, and the presence of the AMR export pump YbtPQ, may lead to the development of non-carbapenemase producing (NCP) carbapenem resistant strains, which may become increasingly difficult to treat. Surveillance of pESBL-PH-2018-like plasmids is recommended to monitor further plasmid dissemination; a critical task as current surveillance efforts mainly consider clonal expansion of resistance encoding strains.

## Data Availability

Long and short read data for strain UHD-2018, including pESBL-PH-2018, can be found on the Sequence Read Archive (SRA) on NCBI under the Bioproject accession number: PRJNA1150551, https://www.ncbi.nlm.nih.gov/sra/?term=PRJNA1150551. UHD-2018 chromosome accession: CP168069.1, plasmid pESBL-PH-2018 accession: CP168070.1.

## Supplementary data description

Supplementary file 1: A list of plasmids similar to pESBL-PH-2018 sourced from the NCBI RefSeq Plasmid Database downloaded on July 16, 2024. For each plasmid, NCBI accession, length, geographical source isolation, host strain/sequence type (ST), isolation source, predicted resistome, replicon, and coverage/identity alongside single nucleotide polymorphisms (SNPs) against pESBL-PH-2018 is recorded.

Supplementary file 2: The strain background of isolates carrying plasmids similar to pESBL-PH-2018 is recorded. For each strain which carried a plasmid similar to pESBL-PH-2018, the K and O-locus of the host chromosome is recorded. Additionally, the *in-silico* predicted resistance, stress, and virulence profile of the host chromosome is recorded. Accessory plasmids, in the host strain, not including pESBL-PH-2018 are recorded and their predicted resistance phenotype is reported.

## References

1. World Health Organisation. Global Priority List of Antibiotic-resistant bacteria to guide research, discovery, and development of new antibiotics. 2017.

2. Lester, R., Musicha, P., Kawaza, K., Langton, J., Mango, J., Mangochi, H., Bakali, W., Pearse, O., Mallewa, J., Denis, B., Bilima, S., Gordon, S. B., Lalloo, D. G., Jewell, C. P., & Feasey, N. A. (2022). Effect of resistance to third-generation cephalosporins on morbidity and mortality from bloodstream infections in Blantyre, Malawi: A prospective cohort study. The Lancet Microbe, 3(12), e922–e930. 10.1016/s2666-5247(22)00282-8

3. Chen, J., Allel, K., Zhuo, C., Luo, W., He, N., Yang, X., Guo, Y., Wang, J., Yao, L., Li, J., Lin, Y., Tu, R., Yakob, L., & Zhuo, C. (2024). Extended-spectrum β-lactamase-Producing *Escherichia coli* and *Klebsiella pneumoniae*: Risk factors and economic burden among patients with bloodstream infections. Risk Management and Healthcare Policy, 17, 375–385. 10.2147/rmhp.s453686

4. Wyres, K. L., Hawkey, J., Hetland, M. A., Fostervold, A., Wick, R. R., Judd, L. M., Hamidian, M., Howden, B. P., Löhr, I. H., & Holt, K. E. (2019). Emergence and rapid global dissemination of CTX-M-15-associated *Klebsiella pneumoniae* strain ST307. Journal of Antimicrobial Chemotherapy, 74(3), 577–581. 10.1093/jac/dky492

5. Sewunet, T., Asrat, D., Woldeamanuel, Y., Ny, S., Westerlund, F., Aseffa, A., & Giske, C. G. (2021). High prevalence of blaCTX-M-15 and nosocomial transmission of hypervirulent epidemic clones of *Klebsiella pneumoniae* at a tertiary hospital in Ethiopia. JAC-Antimicrobial Resistance, 3(1). 10.1093/jacamr/dlab001

6. Poulou, A., Voulgari, E., Vrioni, G., Koumaki, V., Xidopoulos, G., Chatzipantazi, V., Markou, F., & Tsakris, A. (2013). Outbreak caused by an ertapenem-resistant, CTX-M-15-Producing *Klebsiella pneumoniae* sequence type 101 clone carrying an OmpK36 Porin variant. Journal of Clinical Microbiology, 51(10), 3176–3182. 10.1128/jcm.01244-13

7. Zhou, K., Lokate, M., Deurenberg, R. H., Arends, J., Lo-Ten Foe, J., Grundmann, H., Rossen, J. W., & Friedrich, A. W. (2015). Characterization of a CTX-M-15 producing *Klebsiella pneumoniae* outbreak strain assigned to a novel sequence type (1427). Frontiers in Microbiology, 6. 10.3389/fmicb.2015.01250

8. Heiden, S. E., Hübner, N., Bohnert, J. A., Heidecke, C., Kramer, A., Balau, V., Gierer, W., Schaefer, S., Eckmanns, T., Gatermann, S., Eger, E., Guenther, S., Becker, K., & Schaufler, K. (2020). A *Klebsiella pneumoniae* ST307 outbreak clone from Germany demonstrates features of extensive drug resistance, hypermucoviscosity, and enhanced iron acquisition. Genome Medicine, 12(1). 10.1186/s13073-020-00814-6

9. Emeraud, C., Figueiredo, S., Bonnin, R. A., Khecharem, M., Ouzani, S., Leblanc, P., Jousset, A. B., Fortineau, N., Duranteau, J., & Dortet, L. (2021). Outbreak of CTX-M-15 extended-spectrum β-lactamase-Producing *Klebsiella pneumoniae* ST394 in a French intensive care unit dedicated to COVID-19. Pathogens, 10(11), 1426. 10.3390/pathogens10111426

10. Navon-Venezia, S., Kondratyeva, K., & Carattoli, A. (2017). *Klebsiella pneumoniae*: A major worldwide source and shuttle for antibiotic resistance. FEMS Microbiology Reviews, 41(3), 252–275. 10.1093/femsre/fux013

11. Hawkey, J., Wyres, K. L., Judd, L. M., Harshegyi, T., Blakeway, L., Wick, R. R., Jenney, A. W., & Holt, K. E. (2022). ESBL plasmids in *Klebsiella pneumoniae*: Diversity, transmission and contribution to infection burden in the hospital setting. Genome Medicine, 14(1). 10.1186/s13073-022-01103-0

12. van Almsick, V., Schuler, F., Mellmann, A., & Schwierzeck, V. (2022). The use of long-read sequencing technologies in infection control: Horizontal transfer of a *bla*_CTX-M-27_ containing lncFII plasmid in a patient screening sample. Microorganisms, 10(3), 491. 10.3390/microorganisms10030491

13. Dolejska, M., Brhelova, E., Dobiasova, H., Krivdova, J., Jurankova, J., Sevcikova, A., Dubska, L., Literak, I., Cizek, A., Vavrina, M., Kutnikova, L., & Sterba, J. (2012). Dissemination of IncFIIK-type plasmids in multiresistant CTX-M-15-producing Enterobacteriaceae isolates from children in hospital paediatric oncology wards. International Journal of Antimicrobial Agents, 40(6), 510–515.

14. Pedersen, T., Tellevik, M. G., Kommedal, Ø., Lindemann, P. C., Moyo, S. J., Janice, J., Blomberg, B., Samuelsen, Ø., & Langeland, N. (2020). Horizontal plasmid transfer among klebsiella pneumoniae isolates is the key factor for dissemination of extended-spectrum β-lactamases among children in Tanzania. mSphere, 5(4). 10.1128/msphere.00428-20

15. Fordham, S. M., Drobniewski, F., Barrow, M., Hutchings, M., Crowther, K., Richards, D., Bolton, P., Mantzouratou, A., & Sheridan, E. (2024). Genetic analyses of rare ESBL ST628 *Klebsiella pneumoniae* detected during a protracted nosocomial outbreak in the United Kingdom. Microorganisms, 12(5), 883. 10.3390/microorganisms12050883

16. Chen, S. (2023). Ultrafast one-pass FASTQ data preprocessing, quality control, and deduplication using fastp. iMeta, 2(2). 10.1002/imt2.107

17. De Coster, W., D’Hert, S., Schultz, D. T., Cruts, M., & Van Broeckhoven, C. (2018). NanoPack: visualizing and processing long-read sequencing data. Bioinformatics, 34(15), 2666–2669. 10.1093/bioinformatics/bty149

18. Wick, R. R., Judd, L. M., Cerdeira, L. T., Hawkey, J., Méric, G., Vezina, B., Wyres, K. L., & Holt, K. E. (2021). Trycycler: consensus long-read assemblies for bacterial genomes. Genome Biology, 22(1). 10.1186/s13059-021-02483-z

19. Kolmogorov, M., Yuan, J., Lin, Y., & Pevzner, P. A. (2019). Assembly of long, error-prone reads using repeat graphs. Nature Biotechnology, 37(5), 540–546. 10.1038/s41587-019-0072-8

20. Wick, R. R., & Holt, K. E. (2021). Benchmarking of long-read assemblers for prokaryote whole genome sequencing. F1000Research, 8, 2138. 10.12688/f1000research.21782.4

21. Vaser, R., & Šikić, M. (2021). Time- and memory-efficient genome assembly with Raven. Nature Computational Science, 1(5), 332–336. 10.1038/s43588-021-00073-4

22. Bouras, G., Judd, L. M., Edwards, R. A., Vreugde, S., Stinear, T. P., & Wick, R. R. (2024). How low can you go? Short-read polishing of Oxford Nanopore bacterial genome assemblies. Microbial Genomics, 10(6). 10.1099/mgen.0.001254

23. Ondov, B. D., Treangen, T. J., Melsted, P., Mallonee, A. B., Bergman, N. H., Koren, S., & Phillippy, A. M. (2016). Mash: fast genome and metagenome distance estimation using MinHash. Genome Biology, 17(1). 10.1186/s13059-016-0997-x

24. Feldgarden, M., Brover, V., Gonzalez-Escalona, N., Frye, J. G., Haendiges, J., Haft, D. H., Hoffmann, M., Pettengill, J. B., Prasad, A. B., Tillman, G. E., Tyson, G. H., & Klimke, W. (2021). AMRFinderPlus and the reference gene catalog facilitate examination of the genomic links among antimicrobial resistance, stress response, and virulence. Scientific Reports, 11(1). 10.1038/s41598-021-91456-0

25. Schwengers, O., Jelonek, L., Dieckmann, M. A., Beyvers, S., Blom, J., & Goesmann, A. (2021). Bakta: Rapid and standardized annotation of bacterial genomes via alignment-free sequence identification. Microbial Genomics, 7(11). 10.1099/mgen.0.000685

26. Che, Y., Yang, Y., Xu, X., Břinda, K., Polz, M. F., Hanage, W. P., & Zhang, T. (2021). Conjugative plasmids interact with insertion sequences to shape the horizontal transfer of antimicrobial resistance genes. Proceedings of the National Academy of Sciences, 118(6). 10.1073/pnas.2008731118

27. Croucher, N. J., Page, A. J., Connor, T. R., Delaney, A. J., Keane, J. A., Bentley, S. D., Parkhill, J., & Harris, S. R. (2014). Rapid phylogenetic analysis of large samples of recombinant bacterial whole genome sequences using Gubbins. Nucleic Acids Research, 43(3), e15–e15. 10.1093/nar/gku1196

28. Evans, D., Sundermann, A., Griffith, M., Rangachar Srinivasa, V., Mustapha, M., Chen, J., Dubrawski, A., Cooper, V., Harrison, L., & Van Tyne, D. (2023). Empirically derived sequence similarity thresholds to study the genomic epidemiology of plasmids shared among healthcare-associated bacterial pathogens. eBioMedicine, 93, 104681. 10.1016/j.ebiom.2023.104681

29. Page, A. J., Cummins, C. A., Hunt, M., Wong, V. K., Reuter, S., Holden, M. T., Fookes, M., Falush, D., Keane, J. A., & Parkhill, J. (2015). Roary: Rapid large-scale prokaryote pan genome analysis. Bioinformatics, 31(22), 3691–3693. 10.1093/bioinformatics/btv421

30. Alikhan, N., Petty, N. K., Zakour, N. L. B., & Beatson, S. A. (2011). BLAST Ring Image Generator (BRIG): simple prokaryote genome comparisons. BMC Genomics, 12(1). 10.1186/1471-2164-12-402

31. Wong, J. L. C., Romano, M., Kerry, L. E., Kwong, H., Low, W., Brett, S. J., Clements, A., Beis, K., & Frankel, G. (2019). OmpK36-mediated Carbapenem resistance attenuates ST258 *Klebsiella pneumoniae* in vivo. Nature Communications, 10(1). 10.1038/s41467-019-11756-y

32. Negeri, A. A., Mamo, H., Gahlot, D. K., Gurung, J. M., Seyoum, E. T., & Francis, M. S. (2023). Characterization of plasmids carrying *bla*_CTX-M_ genes among extra-intestinal *Escherichia coli* clinical isolates in Ethiopia. Scientific Reports, 13(1). 10.1038/s41598-023-35402-2

33. Wranne, M. S., Karami, N., KK, S., Jaén-Luchoro, D., Yazdanshenas, S., Lin, Y., Kabbinale, A., Flach, C., Westerlund, F., & Åhrén, C. (2024). Author correction: Comparison of CTX-M encoding plasmids present during the early phase of the ESBL pandemic in western Sweden. Scientific Reports, 14(1). 10.1038/s41598-024-67491-y

34. Minja, C. A., Shirima, G., & Mshana, S. E. (2021). Conjugative plasmids disseminating CTX-M-15 among human, animals and the environment in Mwanza Tanzania: A need to intensify one health approach. Antibiotics, 10(7), 836. 10.3390/antibiotics10070836

35. Harmer, C. J., Lebreton, F., Stam, J., McGann, P. T., & Hall, R. M. (2022). Mechanisms of IS *26*-Mediated Amplification of the *aphA1* Gene Leading to Tobramycin Resistance in an *Acinetobacter baumannii* Isolate. Microbiology Spectrum, 10(5). 10.1128/spectrum.02287-22

36. Shropshire, W. C., Konovalova, A., McDaneld, P., Gohel, M., Strope, B., Sahasrabhojane, P., Tran, C. N., Greenberg, D., Kim, J., Zhan, X., Aitken, S., Bhatti, M., Savidge, T. C., Treangen, T. J., Hanson, B. M., Arias, C. A., & Shelburne, S. A. (2022). Systematic analysis of mobile genetic elements mediating β-lactamase gene amplification in noncarbapenemase-producing carbapenem-resistant *Enterobacterales* bloodstream infections. mSystems, 7(5). 10.1128/msystems.00476-22

37. Farzand, R., Rajakumar, K., Barer, M. R., Freestone, P. P., Mukamolova, G. V., Oggioni, M. R., & O’Hare, H. M. (2021). A virulence associated Siderophore importer reduces antimicrobial susceptibility of *Klebsiella pneumoniae*. Frontiers in Microbiology, 12. 10.3389/fmicb.2021.607512

38. Gullberg, E., Albrecht, L. M., Karlsson, C., Sandegren, L., & Andersson, D. I. (2014). Selection of a Multidrug resistance plasmid by sublethal levels of antibiotics and heavy metals. mBio, 5(5). 10.1128/mbio.01918-14

39. Shropshire, W. C., Aitken, S. L., Pifer, R., Kim, J., Bhatti, M. M., Li, X., Kalia, A., Galloway-Peña, J., Sahasrabhojane, P., Arias, C. A., Greenberg, D. E., Hanson, B. M., & Shelburne, S. A. (2020). IS26-mediated amplification of *bla*_OXA-1_ and *bla*_CTX-M-15_ with concurrent outer membrane Porin disruption associated with *de novo* carbapenem resistance in a recurrent bacteraemia cohort. Journal of Antimicrobial Chemotherapy, 76(2), 385–395. 10.1093/jac/dkaa447

40. Straub, J. (2013). An environmental Risk Assessment for Human-Use Trimethoprim in European surface waters. Antibiotics, 2(1), 115–162. 10.3390/antibiotics2010115

41. Fordham, S. M. E., Barrow, M., Mantzouratou, A., & Sheridan, E. (2024). Genomic analyses of an *Escherichia coli* and *Klebsiella pneumoniae* urinary tract co-infection using long-read nanopore sequencing. MicrobiologyOpen, 13(1). 10.1002/mbo3.1396

